# Loss of neutrophil Shp1 produces hemorrhagic and lethal acute lung injury

**DOI:** 10.1101/2024.05.23.595575

**Authors:** SF Moussavi-Harami, SJ Cleary, M Magnen, Y Seo, C Conrad, BC English, L Qiu, KM Wang, CL Abram, CA Lowell, MR Looney

## Abstract

The acute respiratory distress syndrome (ARDS) is associated with significant morbidity and mortality and neutrophils are critical to its pathogenesis. Neutrophil activation is closely regulated by inhibitory tyrosine phosphatases including Src homology region 2 domain containing phosphatase-1 (Shp1). Here, we report that loss of neutrophil Shp1 in mice produced hyperinflammation and lethal pulmonary hemorrhage in sterile inflammation and pathogen-induced models of acute lung injury (ALI) through a Syk kinase-dependent mechanism. We observed large intravascular neutrophil clusters, perivascular inflammation, and excessive neutrophil extracellular traps in neutrophil-specific Shp1 knockout mice suggesting an underlying mechanism for the observed pulmonary hemorrhage. Targeted immunomodulation through the administration of a Shp1 activator (SC43) reduced agonist-induced reactive oxygen species *in vitro* and ameliorated ALI-induced alveolar neutrophilia and NETs *in vivo*. We propose that the pharmacologic activation of Shp1 has the potential to fine-tune neutrophil hyperinflammation that is central to the pathogenesis of ARDS.

## Introduction

The acute respiratory distress syndrome (ARDS), defined as the acute onset of respiratory failure, hypoxemia, and noncardiogenic pulmonary edema, contributes to significant morbidity and mortality across the world (1). Despite advancements in supportive care, mortality associated with ARDS has not improved (1, 2). Failure of therapeutic approaches in earlier ARDS trials is likely secondary to the heterogeneity of the susceptible population as well as the complex and fast-moving pathophysiology of ARDS (3–5). Recently, systemic steroids, anti-IL-6, and JAK-STAT blocking therapies have improved outcomes in patients with COVID-19 ARDS (6–10). The success of these therapies in a homogenous etiology of ARDS further illustrates the potential and need for improved targeted therapies for ARDS and a greater understanding of the syndromal pathogenesis.

Advancements in our understanding of ARDS have long implicated the central role of neutrophils, which contribute to ARDS pathogenesis through the release of intracellular proteases, production of reactive oxygen species (ROS) and formation of neutrophil extracellular traps (NETs) (11–14). Neutrophil activation is regulated by activating tyrosine kinases and inhibitory tyrosine phosphatases including Src homology region 2 domain-containing phosphatase-1 (Shp1, encoded by *PTPN6*) (15–21). Shp1 is a cytosolic protein tyrosine phosphatase (PTP) expressed in hematopoietic cells and to a lesser extent, endothelial cells and epithelial cells. Shp1 is recruited by inhibitory receptors through binding to immunoreceptor tyrosine-based inhibitory motifs (ITIM), and dephosphorylates proteins downstream of cytokine receptors including GM-CSF1R, IL-3R, IL-4R, IL-13R, interferon, and integrins (15). Global deficiency of Shp1 in mouse models, termed *motheaten* (*me*) mice, leads to autoimmunity, inflammatory dermatitis, pneumonitis and death (15, 20–23). Detailed cell-specific knockout experiments have established a critical role for Shp1 in regulating myeloid lineage cells; cell-specific Shp1 knockout in dendritic cells or neutrophils recapitulates aspects of the global loss of this protein in mice (16). In patients, Shp1 mutations are associated with neutrophilic dermatitis and chronic obstructive pulmonary disease (COPD), the latter suggesting a role for Shp1 dysregulation in human lung diseases (24–26). Shp1 has been mainly studied in autoimmunity and its role in acute lung injury (ALI) has not been investigated.

Here, using cell-specific Shp1 deletion, we have established a critical role for neutrophil Shp1 in tissue injury and pulmonary hemorrhage in the setting of sterile inflammation, bacterial (*P. aeruginosa*), and viral (SARS-CoV-2) infections. We observed striking hyperinflammation and lethal pulmonary hemorrhage that was dependent on Syk kinase signaling but independent of canonical peptidyl arginine deiminase 4 (PAD4)-dependent NETosis. Finally, through administration of the Shp1-activating small molecule SC43 (27, 28), we inhibited agonist-induced neutrophil ROS production *in vitro* and reduced alveolar neutrophilia and NETs *in vivo*, conceptually supporting Shp1 activation as a therapeutic approach to fine-tune neutrophil function in ARDS.

## Results

### Loss of neutrophil Shp1 exacerbated LPS-induced lung injury and produced lethal pulmonary hemorrhage

We used the intra-tracheal LPS model in cell-specific knockouts of Shp1 to study its role in acute lung inflammation. To study the role of neutrophil Shp1 in acute lung inflammation, *S100a8-Cre* (also known as *MRP8-Cre*) mice were crossed with *Ptpn6^fl/fl^* mice as previously described (16). Since Shp1 deficient mice can have spontaneous inflammation, including pneumonitis, we first tested the baseline analysis of bronchoalveolar lavage (BAL) from mice at 48 hours after PBS intra-tracheal instillation. We did not observe any differences in neutrophilia or alveolar hemorrhage between *Ptpn6^fl/fl^* and *Ptpn6^fl/fl^ S100a8-Cre* mice after PBS instillation (Supplemental Figure 1A-B). However, with intra-tracheal LPS challenge, we observed gross pulmonary hemorrhage on examination of lung tissue and BAL, and present in histologic sections in the *Ptpn6*^fl/fl^ *S100a8*-Cre mice (Figure 1A-E, J-K). In concert with the observed pulmonary hemorrhage, there was increased alveolar neutrophilia (Figure 1F), vascular permeability (Figure 1G), BAL NETs (Figure 1H), and increased mortality (Figure 1I), but similar peripheral blood counts (Supplemental Figure 1C-D). Increased NETs were visually confirmed through intravital lung imaging using the extracellular DNA-labelling dye, Sytox Green (Figure 1L-M). Inflammation induced thrombocytopenia and coagulopathy can lead to *in situ* pulmonary hemorrhage (29). We observed no difference in the peripheral platelet counts in *S100a8-Cre Ptpn6^fl/fl^* mice compared to *Ptpn6^fl/fl^* controls with LPS challenge (Supplemental Figure 1E).

**Figure 1.**
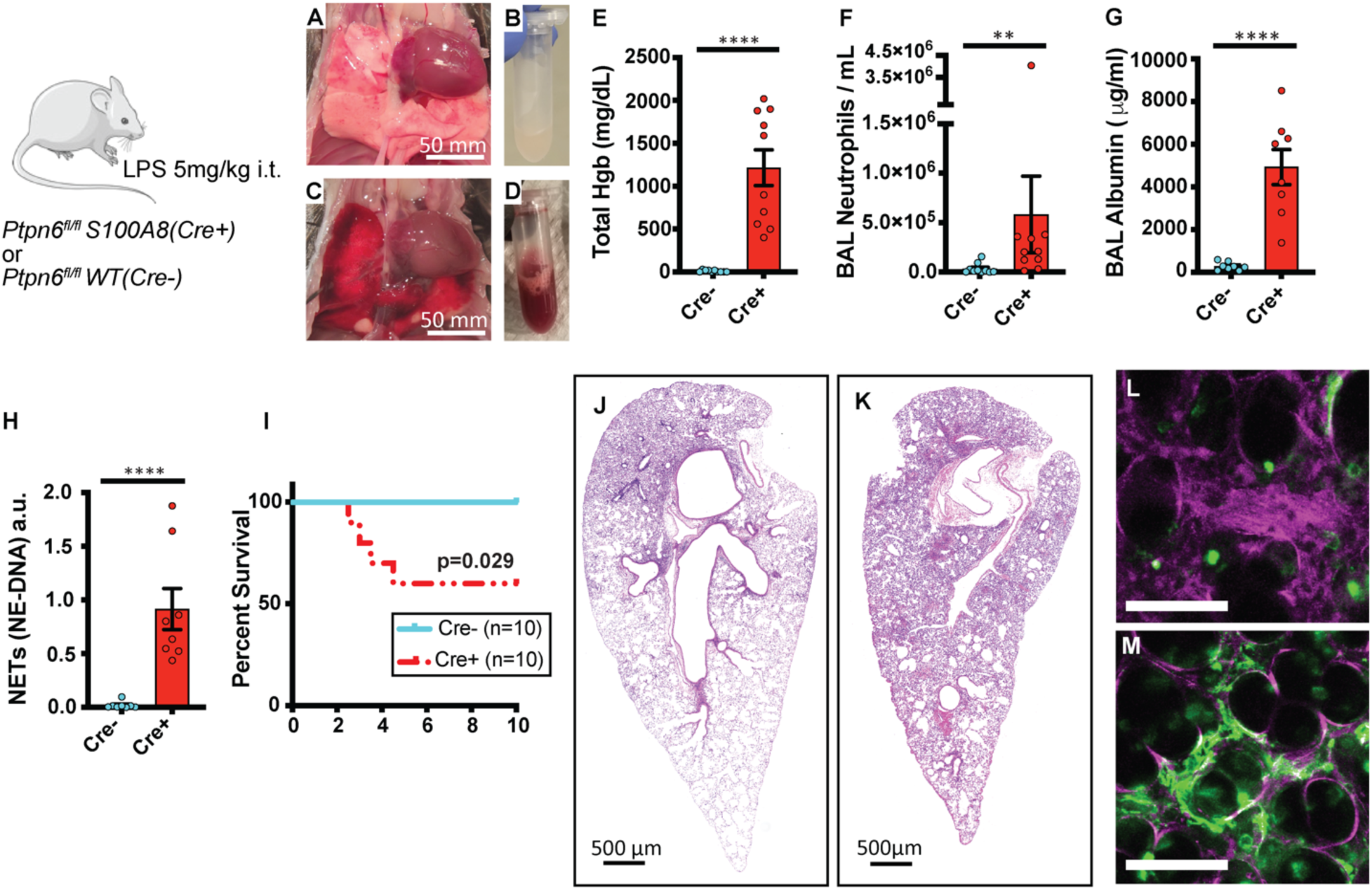
Shp1 deletion in neutrophils leads to severe pulmonary hemorrhage and increased inflammation after LPS-induced lung injury. (A-D) Gross lung and bronchoalveolar lavage (BAL) findings after intra-tracheal LPS in (A,B) *Ptpn6^fl/fl^* and (C,D) *Ptpn6^fl/fl^ S100A8(Cre+)* mice. Quantitative analysis of BAL indicates (E) alveolar hemorrhage, (F) alveolar neutrophilia, (G) increased vascular permeability, and (H) increased BAL NETs in *Ptpn6^fl/fl^ S100A8(Cre+)* mice compared to *Ptpn6^fl/fl^*. (I) Decreased survival in *Ptpn6^fl/fl^ S100A8(Cre+)* mice after LPS. (J-K) H&E staining of lung from (J) *Ptpn6^fl/fl^*and (K) *Ptpn6^fl/fl^ S100A8(Cre+)* mice showing increased inflammation and alveolar hemorrhage after LPS with the loss of Shp1 in neutrophils. (L, M) Intravital image after LPS challenge with Evans Blue (plasma stain) and Sytox Green (NET stain) in (L) *Ptpn6^fl/fl^* and (M) *Ptpn6^fl/fl^ S100A8(Cre+)* indicating exacerbated vascular leak and NETs in *Ptpn6^fl/fl^ S100A8(Cre+)* mice. White bar = 50 µm. *P* values are from unpaired two-tailed t-tests on log10-transformed data (E-H) and log-rank test (I). *\*\*p<0.01, ****p<0.0001*.

Alveolar macrophages are key innate immune cells that serve to recruit neutrophils in the setting of inflammation and infection. To assess the role of alveolar macrophage and dendritic cell Shp1 in ALI, we crossed *Itgax-Cre* (also known as *Cd11c-Cre*) mice with *Ptpn6^fl/fl^* mice as previously described (16). BAL collected 48 hours after intra-tracheal LPS instillation showed no difference in alveolar neutrophilia, alveolar hemorrhage, or neutrophil extracellular traps (NETs) between *Ptpn6^fl/fl^ Itgax-Cre* and *Ptpn6^fl/fl^*mice (Supplemental Figure 2).

### Loss of neutrophil Shp1 led to large intravascular neutrophil clusters and perivascular inflammation

To understand the underlying process leading to the pulmonary hemorrhage that occurs with neutrophils that lack Shp1, we used immunofluorescence imaging and intravital lung imaging to assess neutrophil activity *in vivo* after intra-tracheal LPS instillation, and neutrophil specific functional assays *in vitro*. With Shp1 deletion, we observed an increase in the number of large (volume greater than 5000 µm^3^) intravascular neutrophil clusters partially obstructing the pulmonary arterioles (Figure 2A-G, Supplemental Figures 3& 4, Supplemental Video 1). Staining of red blood cells indicated diffuse alveolar bleeding which increased near pulmonary arterioles (Figure 2D). Furthermore, we observed an increase in alveolar flooding of plasma proteins by intravital imaging following the intravenous administration of Evans blue dye (Figure 2E). *In vitro*, Shp1 deletion led to increased neutrophil ROS production in response to the agonists fMLP and LPS that was dependent on Syk kinase (Supplemental Figure 5A-B). We also observed increased phagocytosis of pH-rhodamine-containing zymosan particles by Shp1-deficient neutrophils (Supplemental Figure 5C-D). These *in vitro* results support our hypothesis that the increased lung inflammation *in vivo* was driven by hyperactive Shp1-deficient neutrophils.

**Figure 2.**
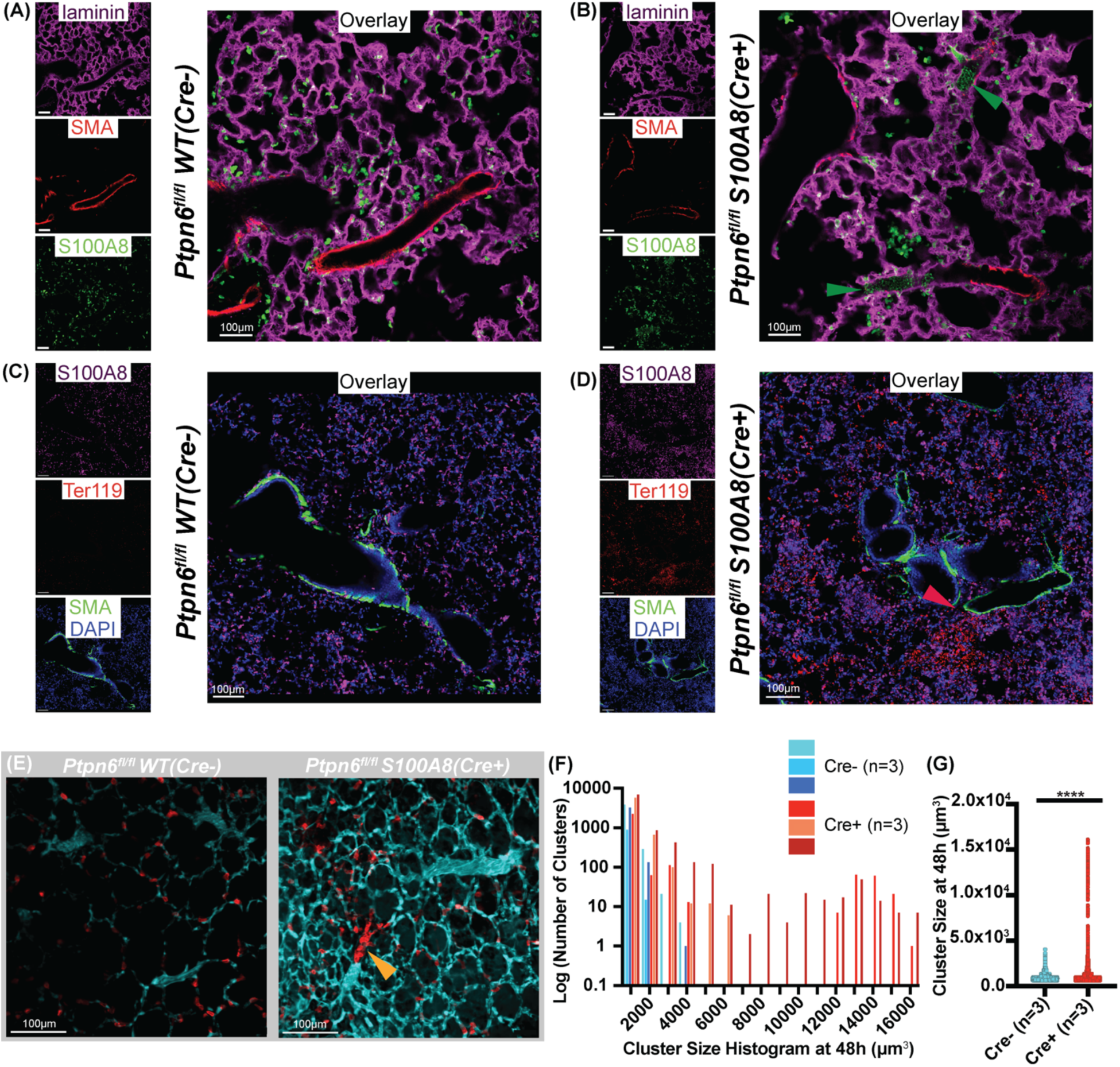
Intravascular neutrophil clusters and perivascular inflammation with loss of neutrophil Shp1. (**A-D**) Immunofluorescence imaging of fixed lung tissue with staining for S100A8 (neutrophils), Ter119 (red blood cells), laminin, and smooth muscle actin (SMA) at 48 hours after LPS challenge from (**A,C**) *Ptpn6^fl/f^* and (**B,D**) *Ptpn6^fl/fl^ S100A8(Cre+)* indicating (**B**) intravascular neutrophil clusters (green arrowheads) and (**D**) perivascular inflammation with preferential alveolar hemorrhage near pulmonary arterioles (red arrowhead). Live intravital lung imaging of (**E**) *Ptpn6^fl/fl^* and *Ptpn6^fl/fl^ S100A8(Cre+)* with anti-Ly6G antibody (neutrophils, red) and plasma albumin labeled by Evan’s blue (cyan) with a large intravascular neutrophil cluster (yellow arrowhead) 48 hours after LPS challenge. (**F**) Histogram of neutrophil cluster size observed over 15 minutes of intravital imaging 48 hours after LPS challenge indicating presence of large clusters. (**G**) Increase in neutrophil cluster size between *Ptpn6^fl/fl^* and *Ptpn6^fl/fl^ S100A8(Cre+)* 48 hours after LPS challenge using the Mann-Whitney non-parametric test. *\*\*\*\*p<0.0001*.

### Shp1 deletion in neutrophils led to pulmonary hemorrhage after bacterial-induced acute lung injury

To test the importance of neutrophil Shp1 in the regulation of ALI caused by a clinically relevant lung infection, mice were challenged with *Pseudomonas aeruginosa* (strain PA103). At 24 hours post-infection, mice lacking neutrophil Shp1 had increased pulmonary hemorrhage (Figure 3A), increased BAL NETs (Figure 3D), worse 3-day survival (Figure 3H) vs. control mice, and impaired bacterial clearance in these mice led to increased extrapulmonary infection and bacteremia (Figure 3E-G). There was no difference in BAL neutrophilia or vascular leak between mice lacking neutrophil Shp1 and controls (Figure 3B-C). Lung histology illustrated increased peri-vascular inflammation and alveolar hemorrhage in neutrophil specific Shp1 knockout mice in comparison to controls (Figure 3I-L).

**Figure 3.**
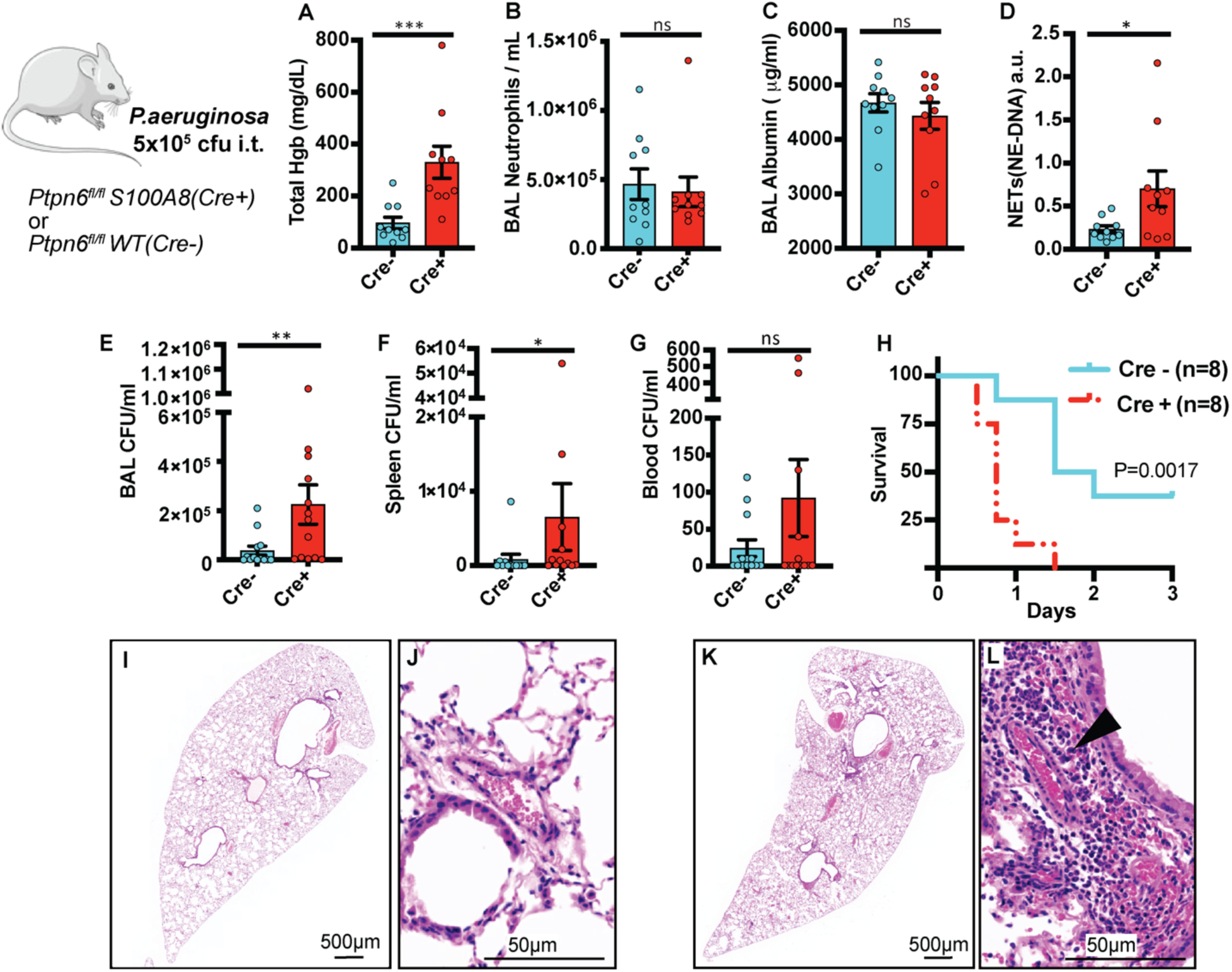
Shp1 deletion in neutrophils leads to a disorganized innate immune response, alveolar hemorrhage, and impaired bacterial clearance after *P. aeruginosa* infection. (**A**) Pulmonary hemorrhage in *Ptpn6^fl/fl^ S100A8(Cre+)* in comparison to *Ptpn6^fl/fl^*, with similar (**B**) alveolar neutrophilia and (**C**) alveolar protein leak but increased (**D**) BAL NETs in *Ptpn6^fl/fl^ S100A8(Cre+)*. (**E**) Increased BAL bacteria, (**F**) spleen bacteria and (**G**) bacteremia in *Ptpn6^fl/fl^ S100A8(Cre+)* with associated (**H**) decreased survival. H&E stained lung tissue after *P. aeruginosa* induced ALI from (**I,J**) *Ptpn6^fl/fl^* and (**K,L**) *Ptpn6^fl/fl^ S100A8(Cre+)* mice with perivascular inflammation (black arrowhead). *P* values are from unpaired two-tailed t-tests on log_10_-transformed data (A-G) and log-rank test (H). *\*p<0.05, **p<0.01, ***p<0.001*.

### Shp1 deletion in neutrophils leads to pulmonary hemorrhage and hyperinflammation in SARS-CoV-2 induced acute lung injury

We also modeled pandemic ARDS with SARS-CoV-2 infections in mice with neutrophil Shp1 deletion. Due to the low affinity of the mouse ACE2 receptor for the viral spike protein, wild-type mice are resistant to infection by ancestral SARS-CoV-2 (30, 31). Recently, mouse-adapted SARS-CoV-2 strains, including MA10, have been developed that better recapitulate human infection (32, 33). To further characterize the role of neutrophil Shp1 in viral respiratory infection, neutrophil specific Shp1 knockout mice and *Ptpn6^fl/fl^* controls were infected with MA10 and monitored for 6 days post-infection. Loss of neutrophil Shp1 resulted in more weight loss, pulmonary hemorrhage, alveolar inflammation, and NETs (Figure 4A-D, F-I), but alveolar protein leak was not significantly increased compared to controls 6 days post-infection (Figure 4E). Overall, these results confirm the critical role of neutrophil Shp1 in regulating inflammation and pulmonary hemorrhage in both bacterial and viral infections.

**Figure 4.**
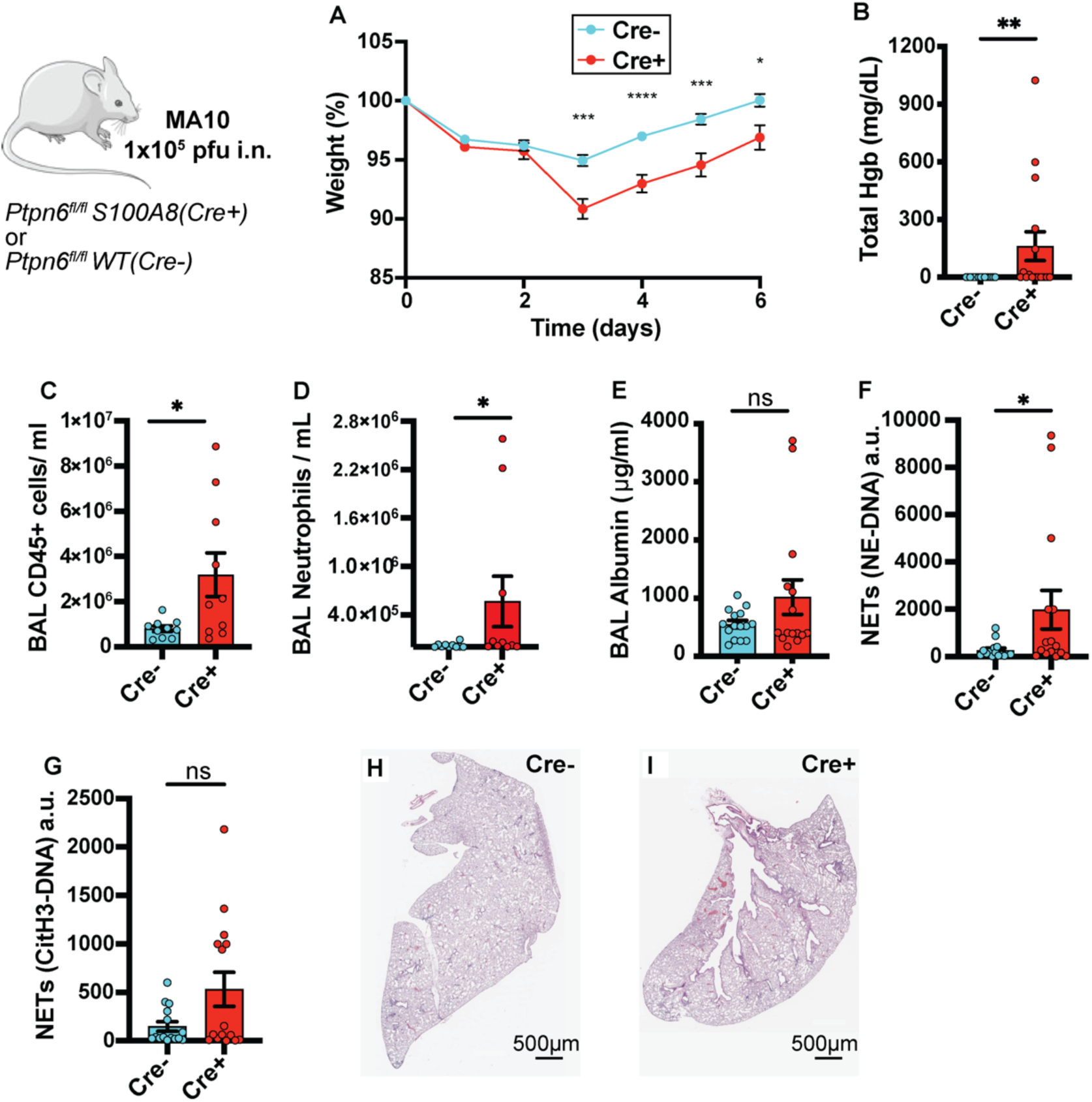
Deletion of Shp1 in neutrophils produces more severe lung injury after SARS-CoV-2 (MA-10) infection. MA-10 infection in *Ptpn6^fl/fl^ S100A8(Cre+)* mice produces increased (**A**) weight loss, (**B**) alveolar hemorrhage, (**C**) alveolar inflammation, (**D**) alveolar neutrophilia, (**E**) similar protein leak, (**F**) increased NE-DNA complexes, and (**G**) similar CitH3-DNA complexes. H&E stained lung tissue from (**H**) *Ptpn6^fl/fl^*(control) and **(I**) *Ptpn6^fl/fl^ S100A8(Cre+)* indicating increased peri-bronchial inflammation. *P* values are from unpaired two-tailed t-tests (A), and unpaired two-tailed t-tests on log_10_-transformed data (B-G). *\*p<0.05, **p<0.01, ***p<0.001, ****p<0.0001*.

### Pulmonary hemorrhage caused by loss of Shp1 was dependent on Syk kinase

Shp1 is required for integrin signaling, and loss of Shp1 in neutrophils leads to hyperadhesion and reduced migration through a Syk kinase-dependent signaling mechanism (16). We hypothesized that loss of Syk kinase in neutrophils would prevent the pulmonary hemorrhage and hyperinflammation observed in *Ptpn6^fl/fl^ S100a8-Cre* mice. We crossed *Syk^fl/fl^* mice with *Ptpn6^fl/fl^ S100a8-Cre* to generate mice with neutrophils lacking both Shp1 and Syk and challenged them with intratracheal LPS. *Syk* kinase deletion in neutrophils that also lacked Shp1 reversed the LPS-induced pulmonary hemorrhage, alveolar neutrophilia, hyperinflammation, and increased NETs, suggesting that Syk kinase signaling is required for the observed phenotype (Figure 5A-I). NETs have been suggested to contribute to pulmonary hemorrhage in trauma and vasculitis and PAD4-dependent histone citrullination is critical for NETosis (34, 35). To better understand the role of NETs in the pulmonary hemorrhage we observed as a result of neutrophil Shp1 deletion, *PAD4^-/-^*mice were crossed to *Ptpn6^fl/fl^ S100a8-Cre,* and challenged with intra-tracheal LPS. As expected, CitH3-DNA NET levels were significantly reduced in *PAD4^-/-^* mice (Supplemental Figure 6E), but there was no change in alveolar hemorrhage, neutrophilia, protein leak, or histologic lung injury (Supplemental Figure 6A-C, F-K). In addition, there was a persistent increase in NE-DNA complex NETs (Supplemental Figure 6D), suggesting that PAD4-independent NET formation was dominant in this model.

**Figure 5.**
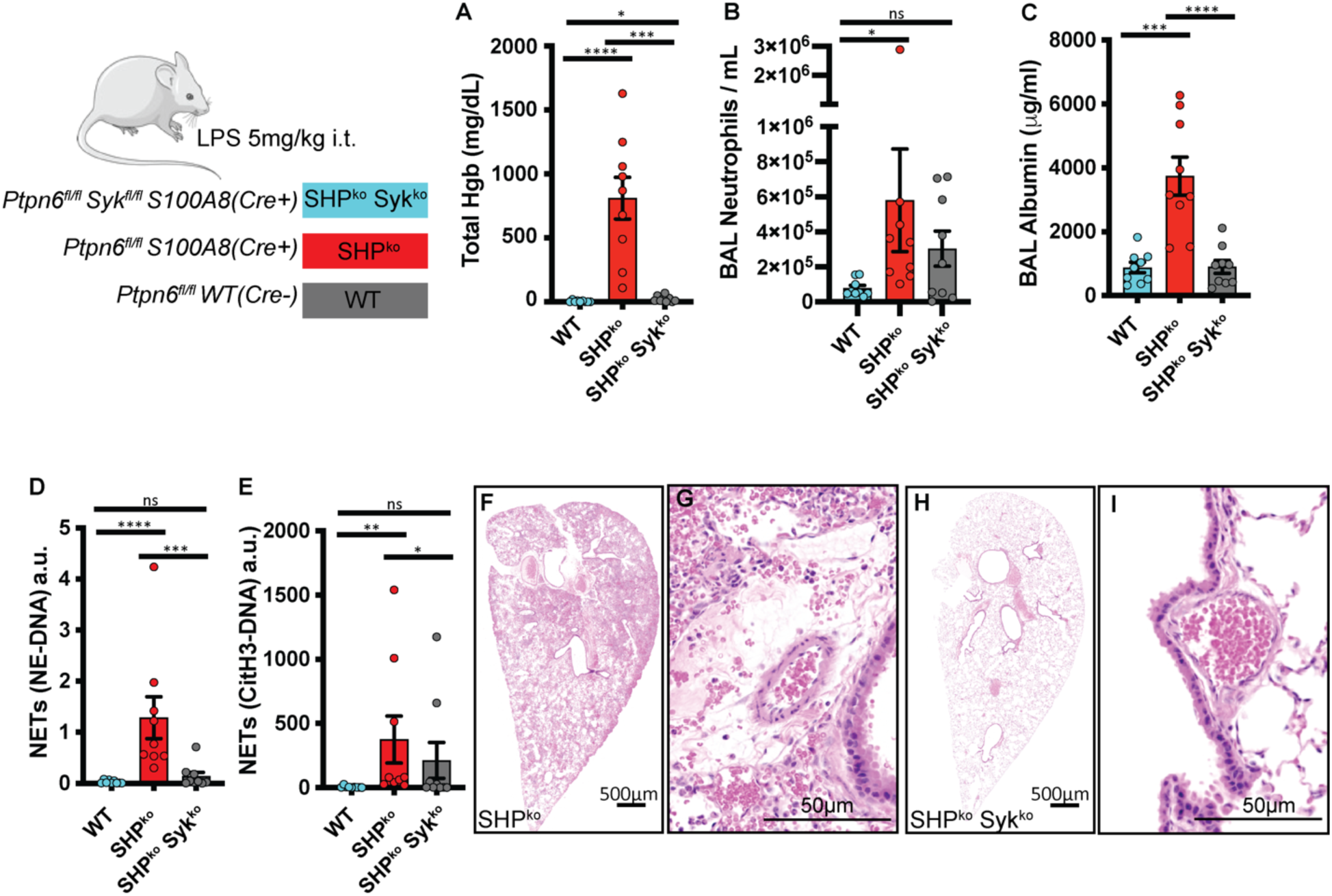
Lung injury from neutrophil Shp1 deletion is dependent on *Syk* kinase signaling. (**A**) Alveolar hemorrhage, (**B**) neutrophilia, (**C**) increased vascular permeability, (**D-E**) increased NE-DNA and CitH3-DNA complexes (NETs) is dependent on *Syk* kinase signaling, H&E-stained lung tissue in (**F-G**) *Ptpn6^fl/fl^ S100A8(Cre+)*, (**H-I**) *Ptpn6^fl/fl^ Syk^fl/fl^ S100A8(Cre+)* indicating perivascular inflammation is dependent on *Syk* kinase signaling. Log_10_ transformed data were analyzed using one-way ANOVA with Tukey’s test for multiple comparisons (A-E). *\*p<0.05*, *\*\*p<0.01, ***p<0.001, ****p<0.0001*.

### Shp1 activation reduced LPS-induced lung inflammation

Our observation that loss of Shp1 in neutrophils produces increased lung injury in multiple models of ALI led us to hypothesize that activation of Shp1 could dampen this response. The small molecule SC43, a derivative of the kinase inhibitor sorafenib, has been described to activate Shp1 in hepatocellular carcinoma cells, leading to dephosphorylation of downstream signaling proteins, including STAT3 (36, 37). *In vivo*, SC43 treatment ameliorated lung fibrosis in a bleomycin-induced model (27). To test whether increased Shp1 activity would reduce inflammation in the LPS ALI model, we treated mice with SC43 prior to intratracheal LPS challenge. Administration of SC43 resulted in reduced alveolar neutrophilia and CitH3-DNA complexes (Figure 6A and D), with similar vascular leak, NE-DNA complexes, and lung injury by histology (Figure 6B-C, E-H). *In vitro*, treatment of neutrophils with SC43 significantly reduced integrin-dependent ROS production stimulated by LPS (Figure 6I-J). These findings support the use of Shp1 activation to modulate neutrophil responses in inflammation and acute lung injury.

**Figure 6.**
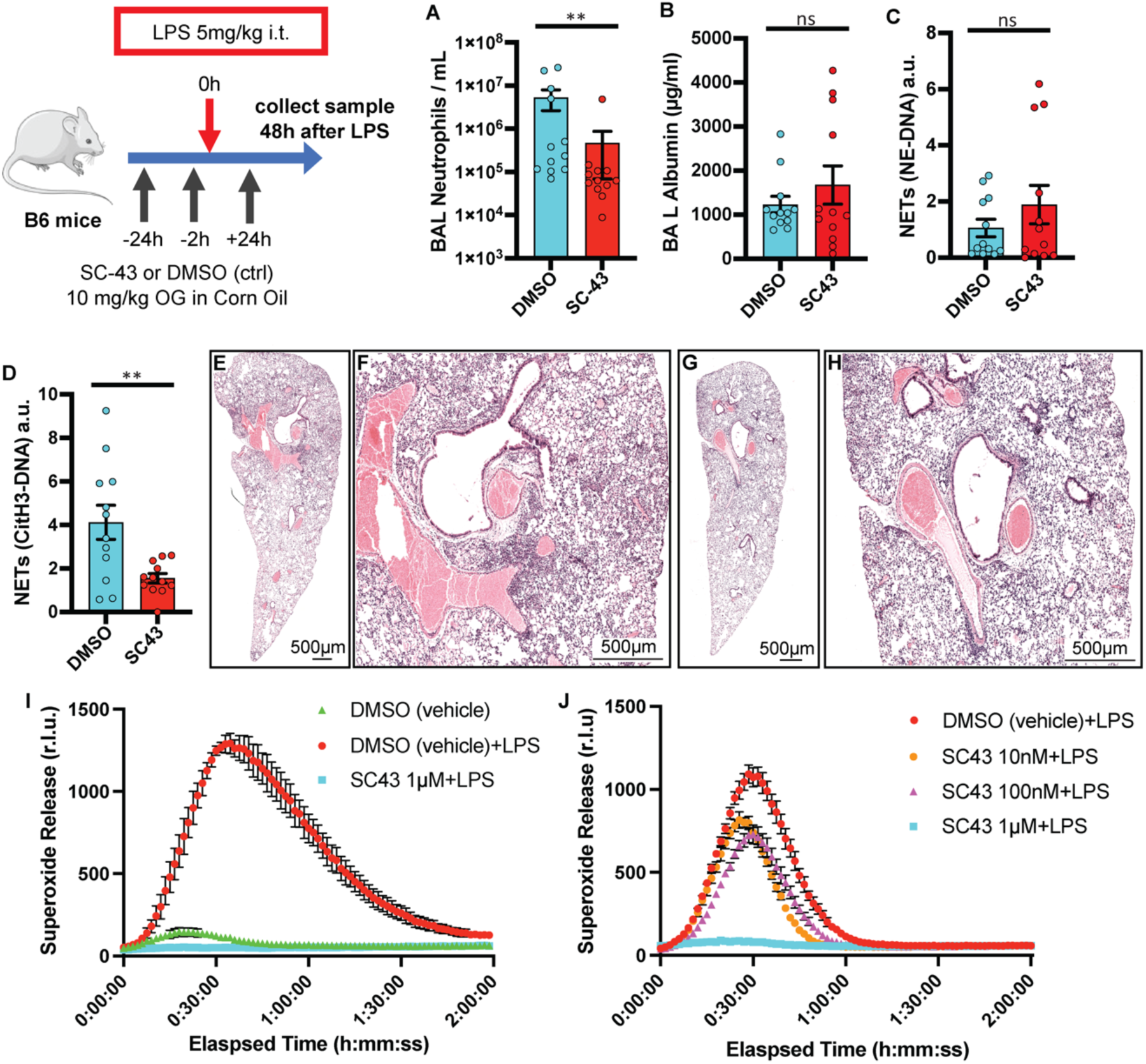
Shp1 activation with SC43 reduces inflammation in LPS-induced lung injury. (**A**) Reduced alveolar neutrophils and (**C**) CitH3-DNA complexes (NETs), with similar (**B**) alveolar protein leak, (**D**) NE-DNA complexes (NETs). Lung H&E stained tissue in (**E,F**) DMSO control and (**G,H**) SC43 treated mice indicating reduced peri-hilar inflammation with SC43 administration. (**I, J**) Dose-dependent reduction in LPS-induced ROS (superoxide release) production with SC43 vs. DMSO (vehicle) control (n=3). *P* values are from unpaired two-tailed t-tests on log_10_-transformed data (A-D). *\*\*p<0.01*.

## Discussion

Excessive neutrophilic inflammation including the release of tissue damaging proteases and extracellular traps is critical to the pathogenesis of ARDS. In response to infection or direct tissue injury, a delicate balance exists between the immune response required for microbial control and tissue damaging hyperinflammation. Prior research into the immunobiology of ARDS has focused on cytokine signaling receptors and immune-activating kinases. However, it is likely that immune dysregulation associated with the loss of inhibitory phosphatases also exacerbates lung injury, as indicated by our observation in mice, where the loss of neutrophil Shp1 led to hyperinflammation and pulmonary hemorrhage in the setting of sterile inflammation, and bacterial and viral infections. Although classically associated with autoimmune lung diseases (38), diffuse alveolar hemorrhage (DAH) may occur as a complication of ARDS (39), and has most recently been observed in a subgroup of patients with SARS-CoV-2 infection with substantial mortality (40). Our experiments established a critical role for neutrophil Shp1 in limiting pulmonary hemorrhage in clinically relevant models of ALI. Using lung histology and intravital imaging, we observed perivascular inflammation and large intravascular neutrophil aggregates in *S100a8-Cre x Ptpn6^fl/fl^*mice challenged with LPS or *P. aeruginosa* suggesting that localized vasculitis may cause the observed pulmonary hemorrhage. Perivascular inflammation was not prominent on histology collected from MA10 infected mice, likely related to the later timepoint utilized in this model of ALI (32). Future work is required to assess tissue level inflammation at earlier times after viral infection.

Syk kinase has previously been shown to be important for neutrophil adhesion and adhesion dependent ROS production (41), therefore we hypothesized that Syk signaling contributes to the observed intravascular neutrophil aggregates and perivascular inflammation. Crossing *Syk^fl/fl^ S100A8-Cre* mice to *Ptpn6^fl/fl^* rescued the observed phenotype *in vivo* and *in vitro* confirming that our neutrophil Shp1 knockout phenotype is dependent on Syk kinase signaling.

Syk inhibitors, including fostamatinib are showing promising results in clinical and animal models of ARDS and our experiments further support these observations (42).

Megakaryocyte/platelet-specific deletion of Shp1 in mice results in decreased platelet responses due to reduced GPVI expression (43). In our current experiments, Shp1 was specifically deleted in neutrophils; hence, platelet function will be intact and normal, and therefore, primary platelet dysfunction is not contributing to the hemorrhage in our model.

NETs contribute to small vessel vasculitis and immunothrombosis associated with ARDS (38). Surprisingly, in the current study, genetic inhibition of histone citrullination, an important molecular trigger for NETosis, did not rescue pulmonary hemorrhage and hyperinflammation associated with the loss neutrophil Shp1. Furthermore, there was persistence of PAD4-independent NETosis as indicated by increased NE-DNA complexes. It is likely that PAD4-independent NET formation is critical in these ALI models and that NETs are involved in destabilizing the lung barrier and leading to hemorrhagic ALI (44). Future work will be required to dissect the role of neutrophil specific proteases including neutrophil elastase in the observed hyperinflammation and pulmonary hemorrhage.

Our therapeutic studies using the Shp1 activator SC43 indicated that it could limit the extent of lung inflammation following LPS administration as observed by reduced alveolar neutrophilia and NET formation. This suggests that Shp1 activation could be a promising pharmacologic approach to fine-tune inflammatory responses. Of note, SC43 will cause global activation of Shp1 including other immune cells, epithelial and endothelial cells. Future studies are required to better dissect the impact of SC43 on the different cell populations present in the alveolus, and of course the development of neutrophil-specific Shp1 activators.

In conclusion, we provide strong evidence that neutrophil Shp1 is critical in limiting hyperinflammation and pulmonary hemorrhage in the setting of ALI. Furthermore, our preclinical studies using an activator of Shp1 support the potential use of Shp1-targeted therapies to reduce neutrophilic responses and limit tissue injury.

## Methods

### Mice

Mice were housed and bred under pathogen-free conditions at the UCSF Laboratory Animal Research Center and all experiments conformed to ethical principles and guidelines approved by UCSF Institutional Animal Care and Use Committee. Age and sex-matched mice at 8 – 12 weeks were used for experimental procedures. All experimental mice were on the C57BL/6J background. C57BL/6J wild-type mice were purchased from Jackson Laboratories. *Itgax-Cre, S100A8-Cre*, *Ptpn6*^ff^, *PAD4^-/-^*mice were used as previously published (16, 45). *Syk^ff^*were used as previously described (16, 41).

### LPS-induced lung injury model

Mice were anesthetized using ketamine i.p. and isoflurane, and intratracheally instilled with LPS (5 µg/g body weight) from *E.coli* (O55:B5, Sigma Aldrich) dissolved in PBS or with sterile PBS for control (46). At the endpoint of LPS challenge, mice were euthanized, and blood was collected from the inferior vena cava. The anterior chest was opened, a vessel clamp was placed on the left main bronchus, and BAL fluid was collected from right lung by inserting a 20-gauge catheter into the trachea through which 500ul of PBS+100µM EDTA was flushed back and forth 3 times. The number of neutrophils in BAL fluid was quantified using flow cytometry using Ly6G and CD11b (gated as shown in Supplemental Figure 7), detection of neutrophil elastase-DNA complexes and CitH3-DNA complexes in BAL fluid was done using ELISA as previously described (45, 47). The left lung was inflated with 4% PFA, and embedded in paraffin for H&E staining, or placed in OCT for frozen section staining.

### Blood counts

Blood was collected via the inferior vena cava into K2EDTA MiniCollect tubes (Greiner) for complete blood counts using a Genesis analyzer (Oxford Science).

### Albumin ELISA

BAL fluid albumin concentration was quantified using Mouse Albumin ELISA Kit (Cat#E99-134, Bethyl Laboratories Inc).

### BAL total hemoglobin

Total hemoglobin was measured using the HemoCue Plasma/Low Hb System (Hemocue®, Brea, CA).

### Lung intravital imaging

Intravital lung microscopy was performed as previously described (45, 48, 49). Following anesthesia with ketamine/xylaxine, tracheostomy was performed, and mice were ventilated with room air plus 1% isoflurane at 125 breaths/min at 10µl/kg body weight tidal volume (Minivent, Harvard Apparatus), with 2-3 cmH2O positive end-expiratory pressure. A thoracic window was inserted into an intercostal incision and a region of the visceral pleura of the left lung was immobilized against the window with 25-30 mmHg negative pressure. Imaging experiments used a Nikon A1r laser scanning confocal microscope (UCSF Biological Imaging Development Center) with a Nikon CFI75 Apo 25XC W objective and excitation using Coherent laser lines (488, 561 and 638 nm). Intravenous injections containing Evans blue (3 mg/kg), Sytox Green (Invitrogen, 0.6 mg/kg), and a non-depleting dose of anti-Ly6G PE antibody (BD Biocciences Clone 1A8, 4 µg per mouse) were given prior to imaging. Imaris Software 9.6 (Oxford Instruments, Oxfordshire, England) was used to detect cell cluster volumes using an automated approach using an initial threshold and size filter to detect an object of interest followed by volume expansion.

### P. aeruginosa-induced lung injury model

Mice were anesthetized with ketamine and isoflurane, PA103 was instilled at 5 x 10^5^ CFU/mouse i.t. to induce pneumonia and lung injury (45, 46). Mice were euthanized at 24 hours and BAL and blood were obtained. Bacterial counts were determined for blood, BAL fluid, or spleen homogenate plated on TSB plates. Colonies present on plates were manually counted after overnight culture at 37°.

### SARS-CoV-2 induced lung injury model

All work was conducted under Biosafety Level 3 (BSL-3) conditions. The following reagent was obtained through BEI Resources, NIAID, NIH: SARS-Related Coronavirus 2, Mouse-Adapted, MA10 Variant (in isolate USA-WA1/2020 backbone), Infectious Clone (ic2019-nCoV MA10) in Calu-3 Cells, NR-55329, contributed by Ralph S. Baric. Vero-TMPRSS2 cells (a gift from Dr. Melanie Ott) were cultured in DMEM supplemented with 10% FBS, penicillin/streptomycin, and L-glutamine (Gibco) in a humidified incubator at 37°C and 5% CO_2_. For propagation, Vero-TMPRSS2 cells were infected with SARS-CoV-2 MA10 in serum-free DMEM and incubated at 37°C, 5% CO_2_. After 72 hours, the supernatant was collected, aliquoted, and stored at -80°C. Viral titer was quantified using a plaque assay in Vero-TMPRSS2 cells. Briefly, 10-fold dilutions of the virus stock were added to Vero-TMPRSS2 cells in a 12-well plate for 1 hour, after which an overlay of 1.2% Avicel RC-581 in DMEM was added. The cells were incubated at 37°C, 5% CO_2_ for 96 hours. The cells were fixed with 10% formalin, stained with crystal violet, and washed with water. The plaques were enumerated to determine the titer of the virus stock. Mouse infections were conducted under ABSL-3 conditions. Mice were anesthetized with i.p. ketamine/xylazine (50 µg/mg / 10 µg/mg) and inoculated with SARS-CoV-2 MA10 via intranasal instillation. Each mouse received a dose of 1 × 10^5^ pfu in a total volume of 40 µL. Infected animals were monitored daily for changes in body weight and clinical signs of illness. At 6 days post-infection, blood and BAL were collected for further analysis.

### Neutrophil functional assays

Bone marrow was harvested in room temperature PBS without calcium and magnesium. Red blood cells were lysed by incubation of bone marrow pellet in 4 ml 0.2% NaCl followed by addition of 10 ml 1.2% NaCl solution, cells were filtered through 70 µm filter and centrifuged at 250 x g. The pellet was resuspended in PBS. Neutrophils were isolated by layering PBS containing bone marrow onto 65% Percoll followed by centrifugation at 440 x g for 30 mins without the brake at room temperature. Isolated cells were resuspended in Hank’s Buffered Salt Solution without calcium or magnesium (Cytiva, Logan, Utah). Adhesion-dependent respiratory burst in the presence of agonists was measured using isoluminol-enhanced chemiluminescence as previously described (16, 50, 51). Neutrophils were stimulated in the presence of fMLP (2 µM) and LPS (5 µg/ml). For the phagocytosis assay, polystyrene flow tubes (Falcon #352054) were coated with PEPTIDE-2000 (pRGD, Sigma Aldrich, Lot SLCJ0890) for 1 hr at room temperature, washed with HBSS (Ca+/Mg+), 3x10^5^ neutrophils were placed per tube with 30µl pH-rhodamine Red zymosan A BioParticles (0.5 mg/ml, Invitrogen, Life Tech, Eugene, OR) and incubated for 2 hrs at 37°C. Samples were labelled with anti-Ly6G-FITC for 30 minutes followed by flow cytometry. Samples were gated for Ly6G positive cells and MFI calculated on PE channel.

### SC43 experiments in LPS-induced ALI model

Wild-type B6 mice were housed in our facility for 2 weeks prior to experiments at 8-12 weeks of age. SC43 (MedChemExpress, Lot 99255) was dissolved in DMSO and diluted in corn oil (Sigma Aldrich) at a final dilution of 10% DMSO or SC43. Mice were dosed with SC43 (10 mg/kg) or DMSO control via gavage needle 24 hours before, 3 hours before, and 24 hours after intra-tracheal LPS challenge. All mice underwent intra-tracheal LPS dosing as above, then BAL and lung tissue were collected 48 hours after LPS challenge.

### Flow cytometry

BAL or bone marrow samples were analyzed using a Fortessa flow cytometer (BD) with gating and measurements using FlowJo software (BD) (representative gating strategies are shown in Supplemental Figure 7). The following anti-mouse antibodies were used for flow cytometry: anti-CD45 BV711 (Biolegend Clone, clone 30-F11, lot B366812); anti-CD3 PE (Biolegend Clone 17A2, lot B263031); anti-Ly6G FITC (BD BioSciences, Clone 1A8, lot 9058981); anti-Ly6C APC-Cy7 (Biolegend Clone HK1.4, lot B309226); anti-Siglec-F BV605 (BD Biosciences Clone E50-2440, lot 1344684); anti-CD11b PE-Cy7 (BD Biosciences Clone M1/70, lot 1179919); anti-CD11c BV421 (Biolegend Clone N418, lot B341930); Live/Dead Fixable Far Red Cell Stain (Invitrogen, Lot 2298166).

### Immunofluorescence imaging

Cryosections were made at 200 µm thickness from lungs fixed by inflation and immersion in 1% formaldehyde in PBS. Sections were prepared and stained as previously described (48, 52). Sections were incubated overnight with antibodies targeting α−smooth muscle actin (SMA) conjugated to FITC (Sigma Aldrich Clone 1A4, Lot 0000196944), S100A8 (R&D systems Clone AF3059, Lot WIT0319021), α−SMA conjugated to Cy3 (Sigma Aldrich Clone 1A4, Lot 00000209582), all at 1:500; and TER119 (Invitrogen Clone TER-119, Lot 1931475) and laminin (Abcam Clone AB11575, Lot 1022401) at 1:250 with 5% normal donkey serum, 0.1% bovine serum albumin and 0.3% triton X-100 in phosphate-buffered saline (PBS). After washing, samples were incubated with Alexa Fluor 488, Cy3, or Alexa Fluor 647-conjugated crossed-absorbed polyclonal secondary antibodies targeting goat, rat or rabbit IgG (Invitrogen Clone A10043 Lot 753765; Invitrogen Clone A11055 Lot 2211210; Invitrogen A21447 Lot 872645; Invitrogen A10043 Lot 753765) at 1:500 in PBS+0.3% triton X-100 overnight. After additional washes, sections were mounted in Vectashield (Vector Laboratories, Cat #H-1700) for standard confocal imaging on a Nikon A1r microscope.

### Histology

Lungs were fixed in 4% (vol/vol) PFA, embedded in paraffin, and stained with H&E by the UCSF Histology and Biomarker Core.

### Statistics

All in vivo and vitro experiments were repeated a minimum of 3 times, unless otherwise noted. To determine significance, 2-tailed Student’s t test was used to compare 2 groups. Skewed results (e.g., BAL hemoglobin) were log-transformed prior to comparison to improve symmetry and normal distribution. Mantel-Cox log-rank test was used for the comparison of survival curves.

All statistical calculations and graphs were generated using GraphPad PRISM. A *P* value less than or eual to 0.05 was considered significant.

## Study Approval

All animal experiments were approved by the Institutional Animal Care and Use Committee at UCSF.

## Author contributions

SFM-H, CAL, and MRL designed the experiments. SFM-H, SJC, MM, BCE conducted experiments and acquired data. SFM-H, CC, KMW, YS, LQ assisted with sample collection and analysis. CAL and CLA provided mice. SFM-H and MM performed statistical analysis. SFM-H, CAL, and MRL wrote the original draft of the manuscript. All authors made contributions to reviewing and editing the manuscript.

## Acknowledgments

We thank the UCSF Biological Imaging Development Colab (BIDC) for assistance with lung intravital imaging as well as the UCSF Parnassus Flow Core supported in part by the NIH Diabetes Research Center grant P30 DK063720. We acknowledge support from NIH grants R35HL161241 and R01AI160167 (to MRL); International Anesthesia Research Society Mentored Research Award (to CC); NIH T32 Research Training in Pediatric Critical Care Medicine 2T32HD049303 and NIH K12 Child Health Research Career Development Program 5K12HD105250 (to SFM-H); and American Society of Transplantation Research Network/CSL Behring Basic Research Grant, AABB Postdoctoral Grant (to SJC). Graphical abstract created with BioRender.com.

## Supplemental materials

Supplemental Movie 1. Intravital lung imaging 48h after LPS instillation with intravascular neutrophil clusters in mice lacking neutrophil Shp1.

