## Supplementary Figure for "Loss of neutrophil Shp1 produces hemorrhagic and lethal acute lung injury"

**Supplementary Data**

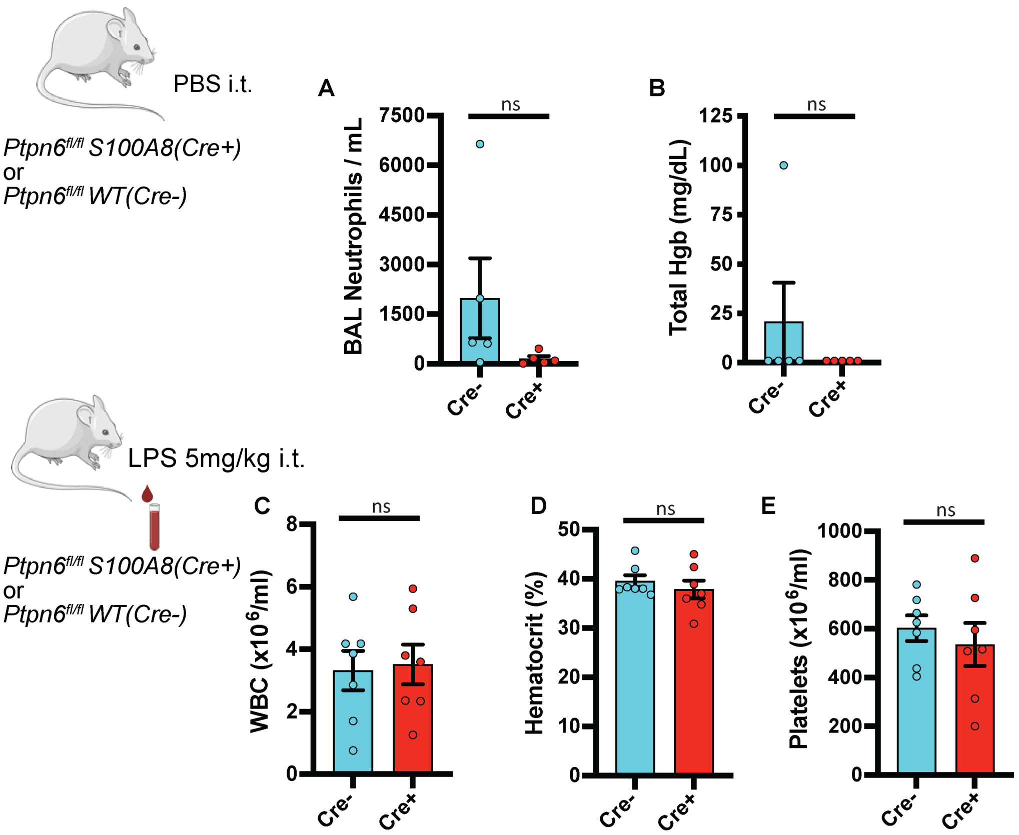


**Supplemental Figure 1. Baseline BAL fluid analysis after PBS instillation and peripheral blood counts after LPS instillation.** Similar (**A**) alveolar inflammation and lack of (**B**) alveolar hemorrhage in *Ptpn6*$^{fl/fl}$ and *Ptpn6*$^{fl/fl}$ *S100A8(Cre+)* 48 hours after intra-tracheal PBS instillation. Similar (**C**) blood WBCs, (**D**) hematocrit, and (**E**) blood platelet counts, 48 hours after intra-tracheal LPS instillation in *Ptpn6*$^{fl/fl}$ and *Ptpn6*$^{fl/fl}$ *S100A8(Cre+)*. Comparison by unpaired two-tailed t-tests on $\log_{10}$-transformed data (A-E).

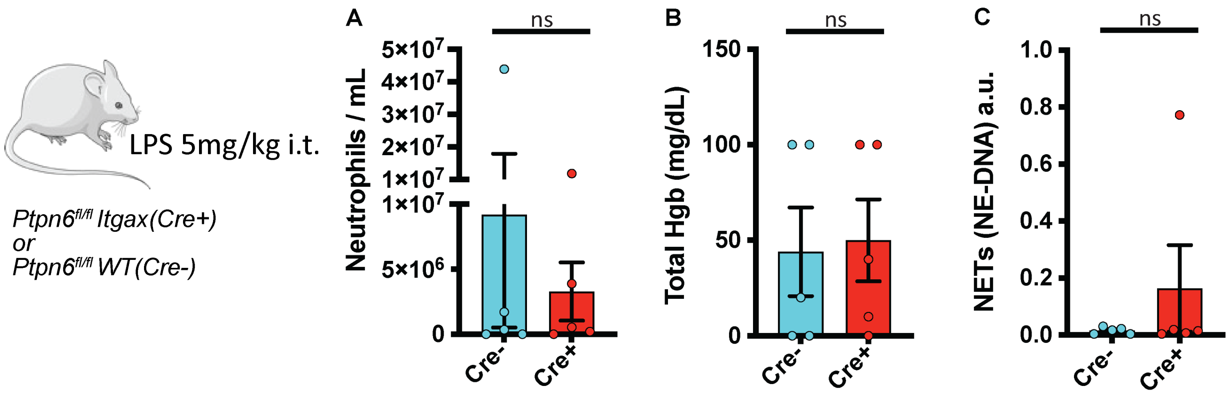


**Supplemental Figure 2. Similar LPS induced lung inflammation with the loss of Shp1 in alveolar macrophages and dendritic cells.** Comparable alveolar (**A**) neutrophilia, (**B**) hemorrhage and (**C**) NETs in *Ptpn6$^{fl/fl}$ Itgax(Cre+)* vs. *Ptpn6$^{fl/fl}$* mice. Comparison by unpaired two-tailed t-tests on $log_{10}$-transformed data (A-C).

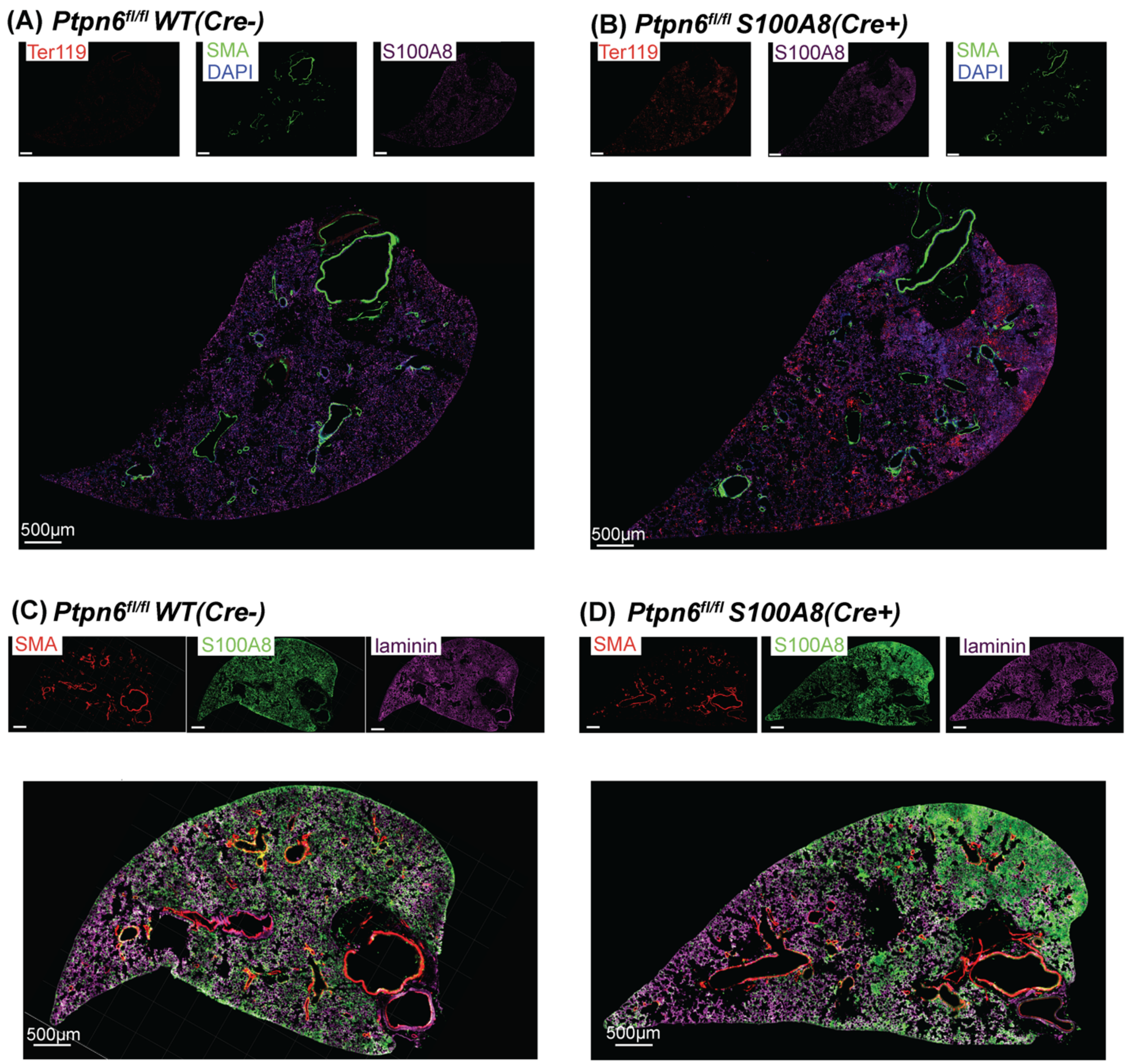


**Supplemental Figure 3. Diffuse neutrophilia and parenchymal hemorrhage with the loss of neutrophil Shp1.** Immunofluorescence imaging of lung tissue with staining for S100A8 (neutrophils), Ter119 (red blood cells), laminin, and smooth muscle actin (SMA) at 48 hours after LPS challenge from (**A**,**C**) *Ptpn6*$^{fl/fl}$ and (**B**,**D**) *Ptpn6*$^{fl/fl}$ *S100A8(Cre+)* and showing increased tissue neutrophils and red blood cell stains in mice with neutrophil Shp1 deletion.

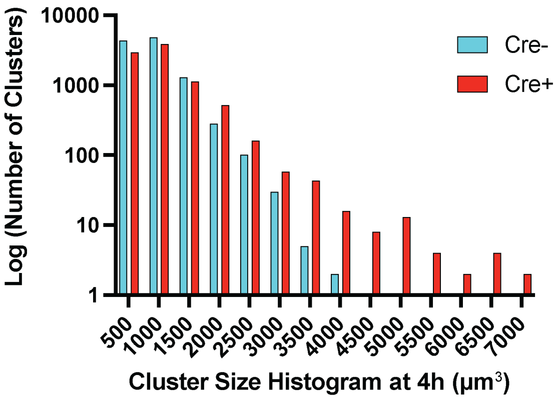


**Supplemental Figure 4. Intravascular neutrophil clusters 4h after intratracheal LPS instillation.** Quantification of intravital lung imaging showing increased numbers and size of intravascular neutrophil clusters 4h after LPS instillation with the loss of Shp1 in neutrophils.

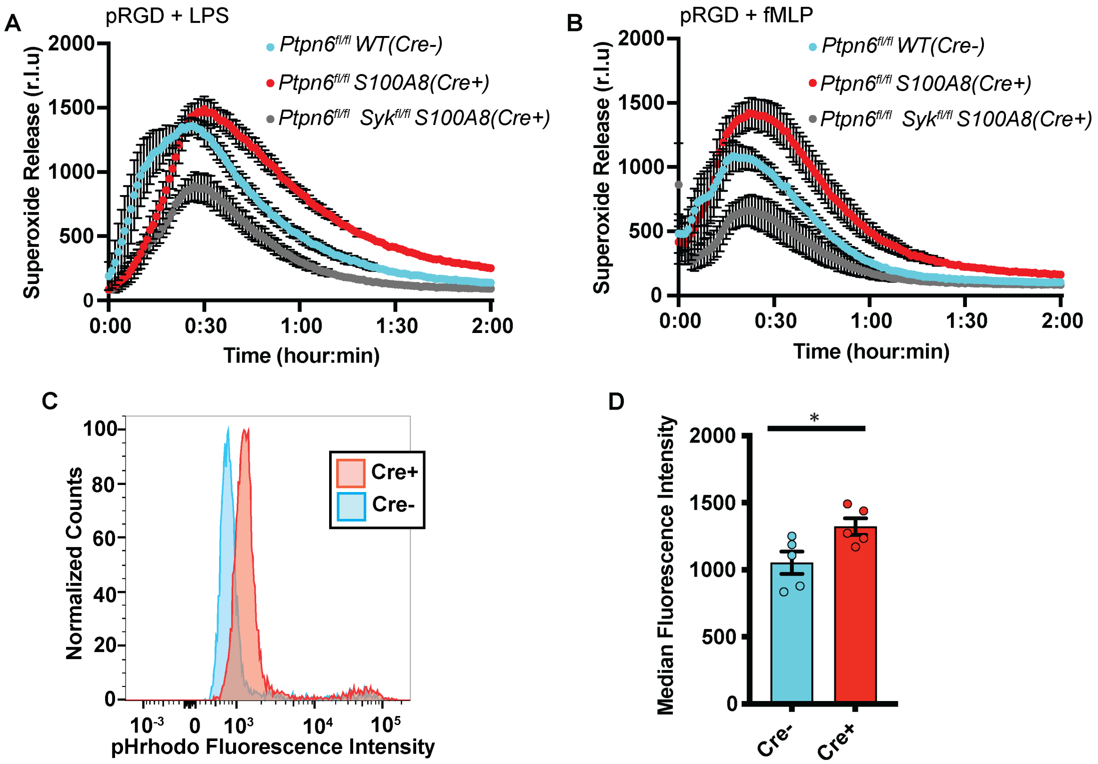


**Supplemental Figure 5. Increased agonist induced reactive oxygen species (ROS) production and increased phagocytosis of pH-Rhodamine-labelled zymosan particles in Shp1 knockout neutrophils.** (**A**) LPS and (**B**) fMLP induced ROS production on RGD-coated surfaces in *Ptpn6$^{fl/fl}$ S100A8(Cre+)* neutrophils is dependent on *Syk* kinase signaling. (**C**) Sample fluorescence curve with (**D**) increased mean fluorescence intensity of pH-Rhodamine zymosan particles indicating increased fluorescence in Shp1 knockout neutrophils (n=5 mice). *P* values are from unpaired two-tailed t-tests on $\log_{10}$-transformed data (F). *$p<0.05$.*

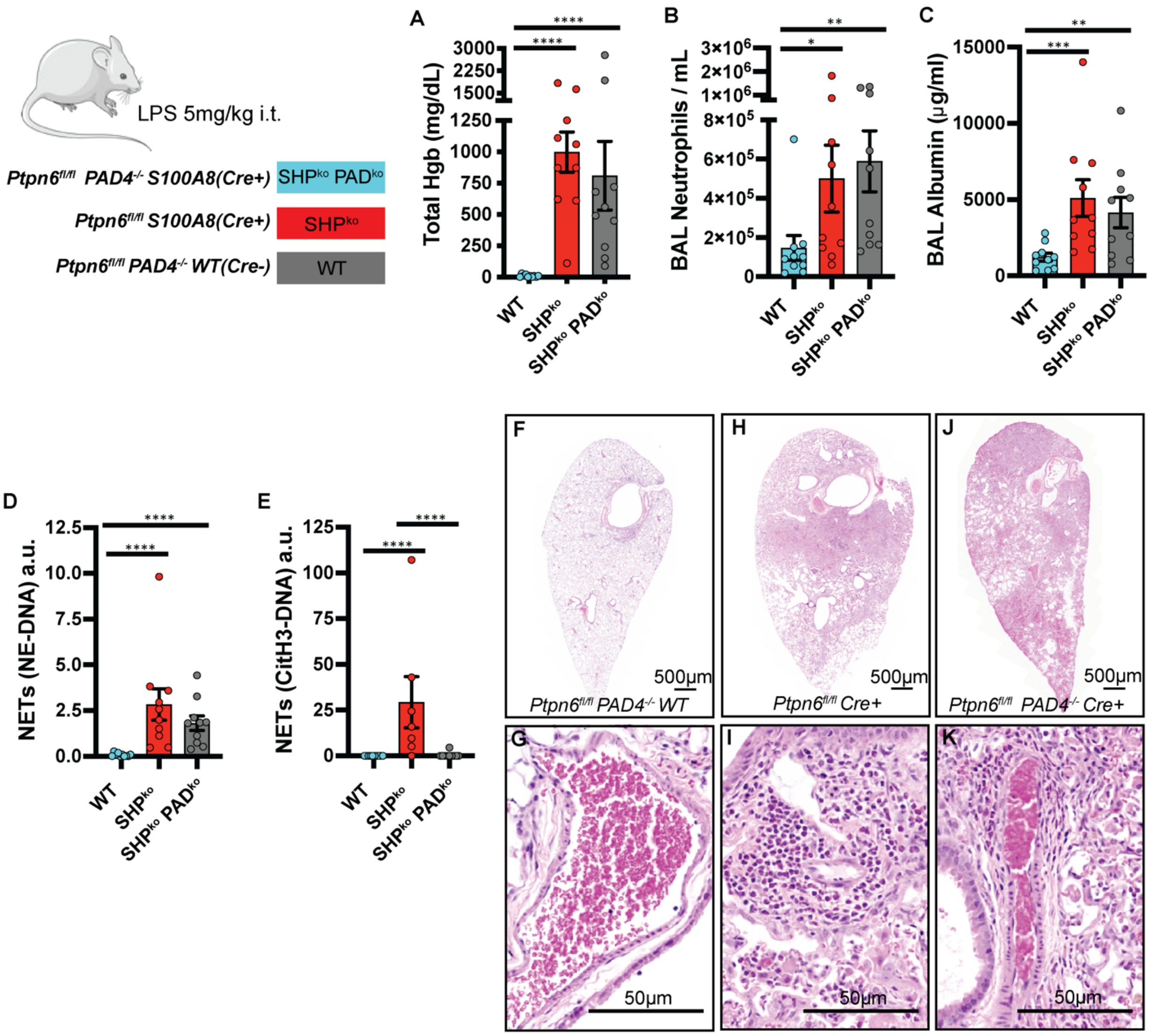


**Supplemental Figure 6. LPS-induced lung hyperinflammation in neutrophil Shp1 knockouts is independent of PAD4 expression.** (**A**) Alveolar hemorrhage, (**B**) BAL neutrophils, (**C**) BAL albumin, (**D**) and BAL NE-DNA NETs are unchanged in *Ptpn6*$^{fl/fl}$ *S100A8(Cre+)* vs. *Ptpn6*$^{fl/fl}$ *PAD4*$^{-/-}$ *S100A8(Cre+)* mice. (**E**) Expected reduction in CitH3-DNA complexes in *Ptpn6*$^{fl/fl}$ *PAD4*$^{-/-}$ *S100A8(Cre+)* mice. (**F-K**) H&E-stained lung tissue in (**F-G**) *Ptpn6*$^{fl/fl}$, (**H-I**) *Ptpn6*$^{fl/fl}$ *S100A8(Cre+)*, (**J-K**) *Ptpn6*$^{fl/fl}$ *PAD4*$^{-/-}$ *S100A8(Cre+)* with similar lung injury in the *Ptpn6*$^{fl/fl}$ *S100A8(Cre+)* and *Ptpn6*$^{fl/fl}$ *PAD4*$^{-/-}$ *S100A8(Cre+)* and (**I,K**) perivascular inflammation. Log$_{10}$ transformed data were analyzed using one-way ANOVA with Tukey's test for multiple comparisons (A-E). ****$p<0.0001$, ***$p<0.001$, **$p<0.01$, *$p<0.05$. Scalebar = 50µm

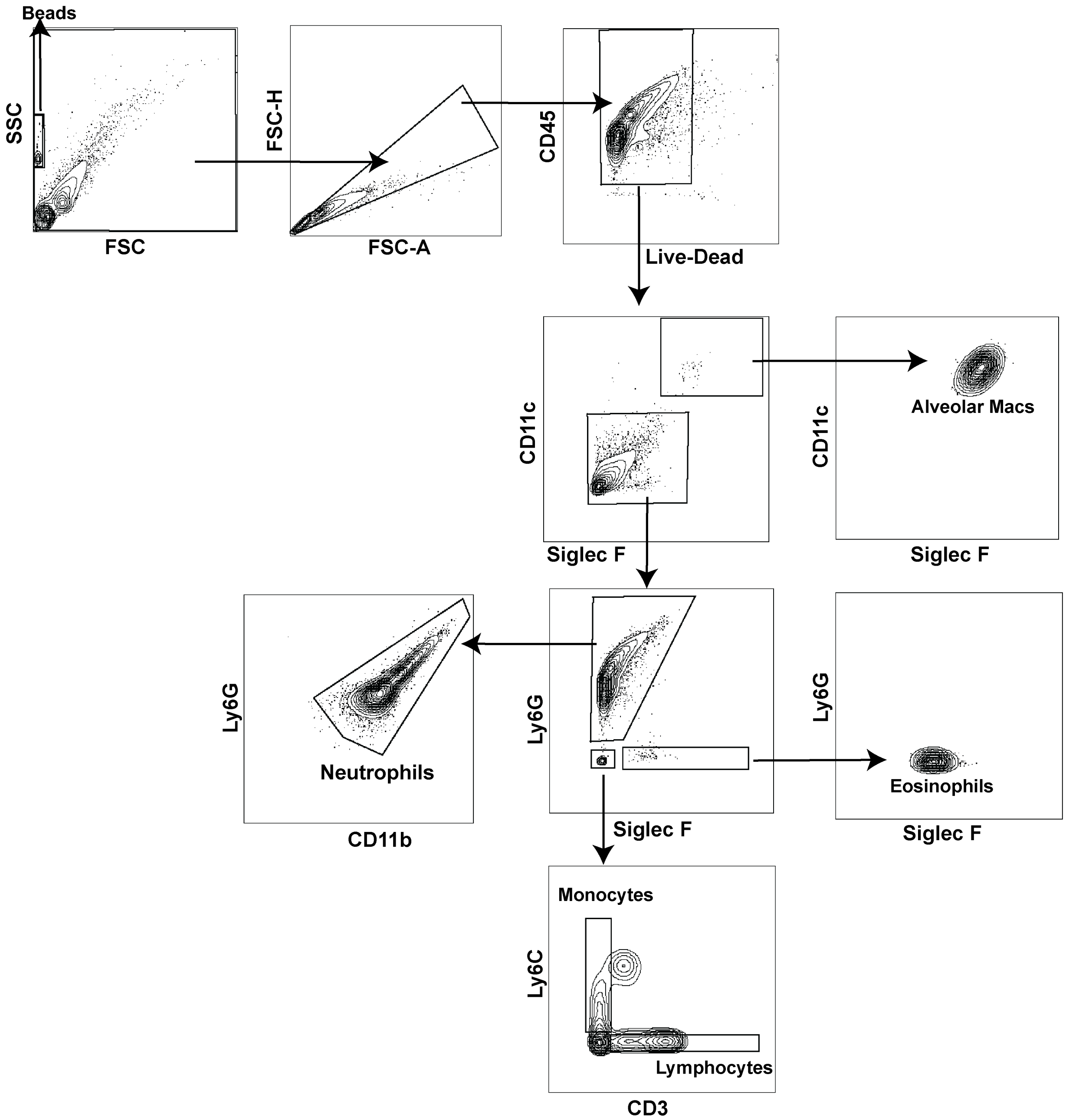


**Supplemental Figure 7.** Sample gating strategy for BAL collected for ALI experiments.
